# ANIMAL-AIDED DESIGN – using a species’ life-cycle to improve open space planning and conservation in cities and elsewhere

**DOI:** 10.1101/150359

**Authors:** Wolfgang W. Weisser, Thomas E. Hauck

## Abstract

Biodiversity underlies many of the ecosystem services demanded by humans. For cities, the design of ‘green infrastructures’ or ‘nature-based solutions’ has been proposed to maintain the provisioning of these services and the preservation of biodiversity. It is unclear, however, how such green infrastructure can be implemented, given existing planning practices that generally ignore biodiversity. Urban open spaces are normally designed by landscape architects with a primary focus on plants, aesthetic design and functionality for human users. As a consequence, conservation of species only plays a minor role, in fact, protected animals are often considered detrimental to the design, e.g. when the need to conserve a protected species demands modifications of a building project. Conversely, conservationists are often in favor of protected areas, also in cities, with little access for humans and no human design.

We propose ‘Animal-Aided Design’ (AAD) as a methodology for the design of urban open spaces, to integrate conservation into open space planning. The basic idea of AAD is to include the presence of animals in the planning process, such that they are an integral part of the design. For AAD, the desired species are chosen at the beginning of a project. The requirements of the target species then not only set boundary conditions for the design, but also serve as an inspiration for the design itself. The aim of AAD is to establish a stable population at the project site, or contribute to population growth of species with larger habitats. AAD thus allows a combination of good urban design with species conservation. We illustrate our approach with designs for urban spaces in Munich.

## INTRODUCTION

Biodiversity is declining worldwide and human land use is the major driver of this decline (Sala, Chapin, Armesto, Berlow, Bloomfield, Dirzo, Huber-Sanwald, Huenneke, Jackson, Kinzig, Leemans, Lodge, Mooney, Oesterheld, Poff, Sykes, Walker Walker and Wall 2000). Because biodiversity underlies many of the ecosystem services that improve human wellbeing, the loss of biodiversity has negative consequences for humans themselves (Millenium Ecosystem Assessment 2005, Cardinale, Duffy, Gonzalez, Hooper, Perrings, Venail, Narwani, Mace, Tilman, Wardle, Kinzig, Daily, Loreau, Grace, Larigauderie, Srivastava and Naeem 2012). To increase the provisioning of ecosystem services and halt the loss of biodiversity, a purely conservation-driven approach, e.g. by protecting intact natural areas, may not be sufficient, because of fast rates of decline. Thus, in addition to the creation of protected areas, biodiversity needs to be supported in areas where humans use the land primarily for their purposes.

Cities are the environment where human appropriation of net primary production and hence impact on the natural environment is greatest. Nevertheless, cities have a role for biodiversity conservation, because of the species living in cities, and because cities nowadays provide the habitat where most humans are most likely to come into contact with wild animal and plants, and profit from the ecosystem services by the species (McKinney 2002, Tanner, Adler, Grimm, Groffman, Levin, Munshi-South, Pataki, Pavao-Zuckerman and Wilson 2014). While species density in cities is lower than in the original habitats that were replaced by cities (Aronson, La Sorte, Nilon, Katti, Goddard, Lepczyk, Warren, Williams, Cilliers and Clarkson 2014, Sol, González-Lagos, Moreira, Maspons and Lapiedra 2014), cities provide an important habitat for a large number of species, from small arthropods to large mammals and birds (Klausnitzer 1991). With increasing urbanization and the increasing loss of urban green space, this biodiversity in cities is, however, declining (McKinney 2001, Grimm, Faeth, Golubiewski, Redman, Wu, Bai and Briggs 2008). Thus, strategies to promote biodiversity in cities are needed (Elmqvist, Fragkias, Goodness, Güneralp, Marcotullio, McDonald, Parnell, Schewenius, Sendstad and Seto 2013). Recent research in urban ecology has begun to unravel features of cities promoting wildlife (Beninde, Veith and Hochkirch 2015, Turrini and Knop 2015), that could be incorporated into planning strategies (Ikin, Le Roux, Rayner, Villaseñor Eyles, Gibbons, Manning and Lindenmayer 2015). In urban and peri-urban areas the incorporation of biodiversity and ecosystem services into planning procedures is often referred to as building a green infrastructure (Benedict and McMahon 2002, European Union 2013, Eggermont, Balian, Azevedo, Beumer, Brodin, Claudet, Fady, Grube, Keune, Lamarque, Reuter, Smith, van Ham, Weisser and Le Roux 2015). The term infrastructure is meant to emphasize the importance of ecosystem services to humans, by making an analogy to technical infrastructures such as roads, bridges, electrical grids etc., that are indispensable for the functioning of a society. Green Infrastructure is often very broadly defined, e.g. as an ‘interconnected network of green space that conserves natural ecosystem values and functions and provides associated benefits to human populations’ (Benedict and McMahon 2002) or even simply as ‘urban and peri-urban green space systems’ (Tzoulas, Korpela, Venn, Yli-Pelkonen, Kaźmierczak, Niemela and James 2007), i.e. without explicit reference to ecosystem services. More recently, ‘nature-based solutions’ have emerged as a competing term that also has several meanings (for a discussion and classification of these concepts see Eggermont, Balian, Azevedo, Beumer, Brodin, Claudet, Fady, Grube, Keune, Lamarque, Reuter, Smith, van Ham, Weisser and Le Roux 2015). Independent of terminology, it is largely unclear how such green infrastructure should be built, in particular how biodiversity conservation and the provision of ecosystem services can be incorporated into particular urban planning projects. Despite frequent calls to integrate biodiversity into urban planning strategies (e.g. Niemelä 1999), the reality is that biodiversity and ecosystem service provision still need to be mainstreamed into landscape and urban planning.

## THE URBAN REALITY – DEVELOPMENT AND CONSERVATION DO NOT FIT TOGETHER

Despite many laudable initiatives such as the creation of ‘green cities’ (Beatley 2012), the more common practice situation in most cities of the world is that there is a conflict between urban planning and the conservation of biodiversity. This conflict primarily arises because biodiversity and ecosystem services play no role during early phases of project planning, when developer, architects, and landscape architects develop their joint project idea. Below, we outline that this exclusion of nature in human design is not a novel phenomenon, but is deeply rooted in human cultural evolution. Biodiversity generally only enters the planning process at a later stage, when the project design has largely been finalized and project specifications and drawings are confronted with environmental laws. Depending on the national legislation, an environmental impact assessment (EIA) may be required, that may also include assessing the impact on biodiversity itself (rather than on other subjects such as water or habitats in general). For example, the Habitats Directive of the European Union (92/43/EEC), adopted 1992, but implemented into the laws of the member states only in the past few years, protects about 400 species listed in Annex IV of the directive, also outside protected areas. Similarly, the Birds Directive (79/409/EEC amended 2009, i.e. 2009/147/EC) protects ca. 500 breeding wild bird species in Europe. For these species, any deliberate capture, killing or disturbance of individuals of the species is forbidden, as well as causing any deterioration or destruction of the breeding sites. In the European Union, therefore, a building project that would result in e.g. the destruction of a breeding site of one of the named species has, in theory, to be stopped or modified to accommodate the requirements of this species. Often, the presence of such a species at a project site is only unraveled during the environmental impact assessment, prompting a conflict between project development and species conservation. Solutions to protect the breeding sites of the species affected, or of the individuals themselves, generally requires some modification of the building project. From the planner’s point of view, such modifications are a nuisance, because of the interference with the original planning intentions, and because they may be costly. In addition, because project development is already advanced, it is often too late to find an acceptable compromise that benefits both humans and biodiversity. More often than not, a minimum solution is sought to fulfill the legal conservation requirements, or the planner directly attempts to obtain an exemption from the law protecting biodiversity. Thus, legal requirements to protect particular species are often a source of conflict in the planning process, rather than a solution that promotes the creation of green infrastructure. The result is often a ‘lose-lose’ situation – the building project is modified resulting in additional costs and potentially inferior design for the developer, and the protected species also suffers from the development, because there is not enough time, money or knowledge to design the compensation measures in a way that the species is not affected or could even benefit from the development.

## DEFENSIVE CONSERVATION AND THE CREATION OF GREEN INFRASTRUCTURE

Importantly, species lists such as the one of the EU Habitat Directive will not protect species that are not on this list. The backdrop of such defensive conservation thus is that it can only preserve the status quo by protecting what is there (cf. Fischman 2006). As a consequence, the legal requirements for protecting biodiversity will not help to create new habitats for species that are not there, at the moment a building project is developed. In cities where competition for space is harsh, biodiversity protection measures will thus not result in new green infrastructure, but can only help to reduce the shrinkage of existing green infrastructure. To be clear: legal biodiversity protection is necessary, in particular in the current situation where biodiversity plays only a minor or no role in most of the urban development. What we argue in this paper however is, that in order to create green infrastructure, biodiversity needs to be mainstreamed into current planning strategies, in order to find solutions that benefit both humans and other species at the same development site. In cities, this is particularly true for those species that can share their habitat with humans, rather than those that live in remnants of their original habitats that were there before the city was built, that are very disturbance-prone and that tolerate little contact with humans. For such species, protected areas within cities without any development are likely to be the best conservation solution. However, with the decline in open green spaces in cities, even these species may benefit from the creation of new habitats that may serve as stepping stones to connect to other protected areas. In this paper, we propose a method that we refer to as Animal-Aided Design, short AAD, that aims to integrate the protection and support of animals into the design of open green spaces, in particular into landscape architectural procedures. The underlying rationale of AAD is that providing habitats for organisms, in particular animals, will only be incorporated into urban open and green space planning, if it is appealing to urban planners and architects, and in particular landscape architects. This can, in our view, only be achieved by developing an approach that takes account of the method of operation in landscape architecture. Importantly AAD is designed to not only protect the biodiversity present at a planning location, but also to create new habitats that would otherwise not be there. Thus, AAD is suited to help creating green infrastructure in urban and peri-urban environments. AAD is a species-centred approach, with all its advantages and disadvantages, as will be discussed below. Before we explain the methodology of AAD in more detail, we will briefly review the different ways in which animals have been traditionally considered in landscape architecture.

## THE HISTORICAL SEPARATION BETWEEN THE ‘HUMAN WORLD’ AND THE ‘ANIMAL WORLD’

The statement that wild animals are part of the urban realm is still alien to most designers, architects and planners. One reason for this is the modern separation of the human world from the natural world, or, seen from a spatial perspective, the separation of city and landscape, into two separated spheres of human life and wildlife, which are considered to be different functional systems. This implicates the exclusion of all kinds of animals – wild animals as well as farm animals – from the urban space (Philo and Wilbert 2000). Similarly, ‘animal-only’ areas or the concept of ‘wilderness’, realized in protected areas and national parks, are in fact representing the same separation of the earth into the human and animal spheres.

This idea of a spatial separation between the human world and wildlife is based on a concept of the relation between human and nature that evolved from the idea of the ‘landscape’ – an idea that was developed as a principle of composition for paintings in the 15th century in Dutch painting workshops (Büttner 2000). In the 16th century, a dissident wing of the British aristocracy and the emerging bourgeoisie started to transfer the pictorial composition principle of landscape into the design of their gardens. The scenery of a painting became the scenery in a landscape garden – i.e. a painting that you could walk into (Hauck 2014, Siegmund 2011). This pictorial principle, that spread when more and more people viewed landscape paintings and visited landscape gardens, became a popular world-view in Europe and the Western world. The so-called picturesque eye was exercised at walks and pleasure trips (Hauck 2014, Trepl 2012, Hussey 1967, Bermingham 1986, Olwig 2002). Animals were an important part of these walk-able paintings and helped to express their symbolic meaning. In the compositional pattern of a landscape, animals became important attributes with different characters – such as sheep, which were, and still are, a crucial part of the image of arcadia, or deer, expressing wilderness (Price 1842). This symbolic relation between the aesthetic character/or expression of a landscape (such as wild, romantic, bucolic, beautiful, etc.) and the character of a species was reified to become a functional, biological or even ecological relation in the 19th century, Animals did not only fit into a specific scenery because of aesthetic or symbolic reasons – but they were a functional and essential part of this landscape. From this follows that to eradicate a species from its indigenous or native landscape meant to damage this landscape aesthetically as well as functionally. On the other hand, the introduction of non-native or alien species disturbs or destructs the balance and harmony of the landscape – functionally as well as aesthetically. In this view, therefore, each landscape has a typical inventory of species, which is fundamental to its aesthetic value and functionality. The parallels to classical traditions of pictorial composition are obvious. All parts of a picture have to merge into a unity, every part has its right place, and no part can be removed or attached without disturbing the harmony of the composition. Animals are part of this composition – in which some species belong into certain landscapes, others do not.

The result of this pictorial understanding of nature is the idea of three spatial relations between man and other species: the first relation is wilderness (Kirchhoff and Trepl 2009), where wild creatures roam around freely and humans are acting as intruders or explorers. The second relation is the city as the civilized sphere of civilized people that are accompanied by pets and (sometimes) vermin; and the third relation is the intermediated sphere of the so-called cultural landscape or Kulturlandschaft, as successor of Arcadia (Trepl 2012), where humans and other species (preferably domesticated animals such as sheep and other peaceful creatures such as songbirds) live harmoniously together. Species that cross the borders between these spheres are often seen as intruders or as abnormal. Thus, different measures are taken to restore the right relations between man and other species – including putting up fences, eradicating animals from certain areas or relocating individuals and populations.

## THE CURRENT GAP BETWEEN LANDSCAPE ARCHITECTURE AND CONSERVATION REFLECTS THE HISTORICAL SCHISM

Currently, landscape architecture does not embrace conservation, and conservation does not normally consider landscape architecture as a means to reach biodiversity targets. Although both professions apply their work to natural environments, they follow fundamentally different objectives. The nature created by landscape architects is dedicated to the recreation purposes of the ‘civilized’ urban human population, thus landscape architecture strives for good design, i.e. controllability, usefulness, aesthetic value and novelty. The creative act of designing something that is novel and manageable is therefore essential for the field. Nature can serve as a source of inspiration, but it is modified and ‘domesticated’ in the landscape architectural design. Classical landscape architecture, with its aim to provide urbanites with a controlled and tamed nature does not care a lot about wild animals or biodiversity when designing urban parks or gardens, apart from the creation of zoos and the application of measures for pest control. In recent years, however, animals increasingly become important pictorial objects in the images made by landscape architects to illustrate their design ideas for urban developments (Fig. 1). For current ‘good’ design, the presence of animals in urban green spaces is important, expressing good ecological order and a harmonious man-nature relationship. However, despite this new inclusion of wild animals into the design of urban green spaces, landscape architecture remains to be the opposite of ‘leaving things as they are in nature’. In contrast, traditional conservation fosters the idea of nature as wilderness, with as little influence of humans as possible, even in the urban environment. As wilderness is the aim, human interference and hence design can only serve to downgrade nature. This fundamental difference in the attitude towards nature still persists, despite reports of an increasing support of native biodiversity by landscape architecture (Müller, Ignatieva, Nilon, Werner and Zipperer 2013).

**Figure 1:**
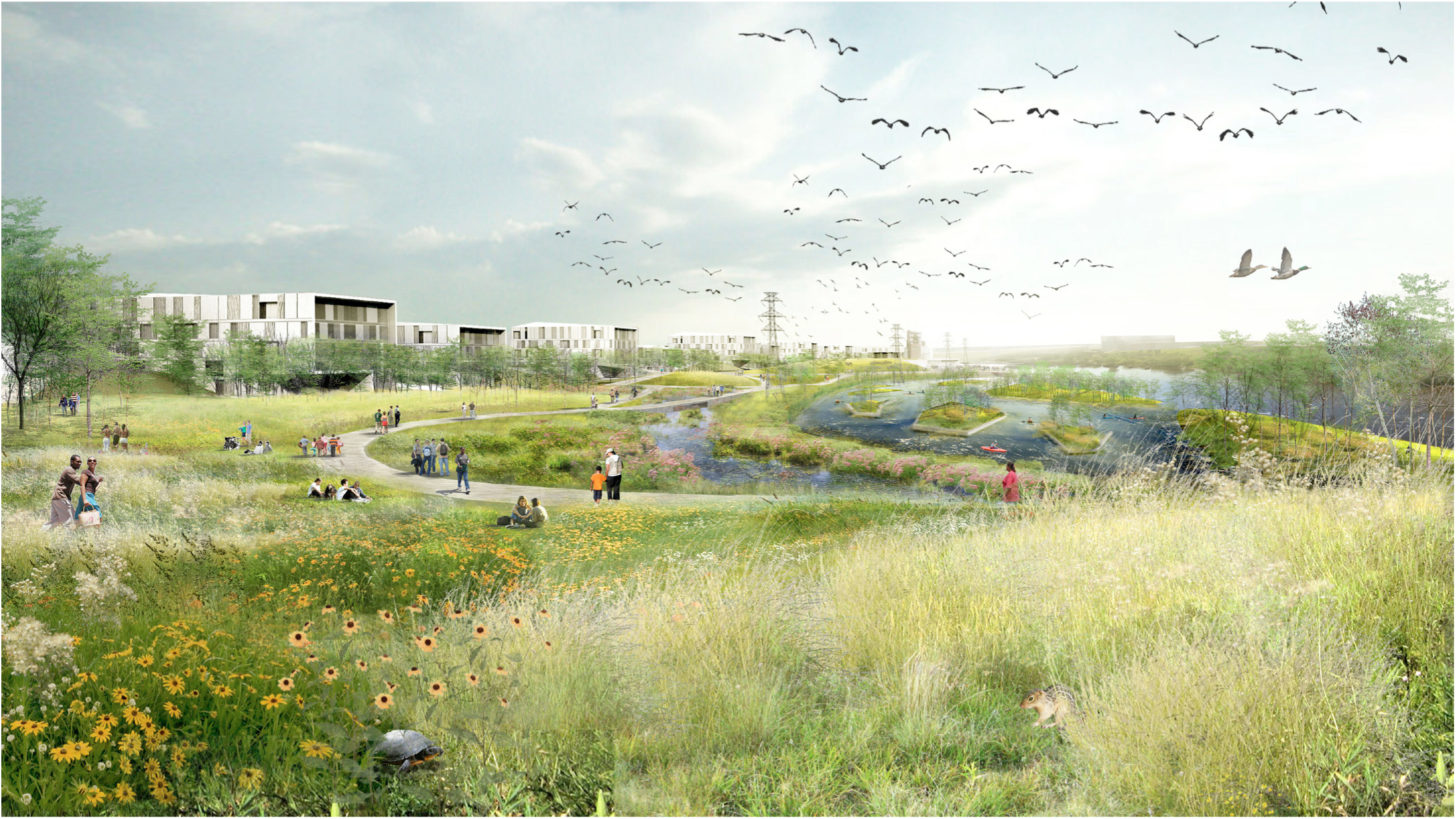
Current landscape architectural designs use the presence of animals as a marker for well-designed urban green spaces. While these animals are not explicitly considered in the landscape architectural design process, they are required to demonstrate the completeness of the design (<u>RiverFirst project by TLS Landscape Architecture</u>). The animals are expected to colonize and live in the designed habitats.

Unfortunately, most of the current calls for cooperation between landscape architecture and conservation are calls to ‘be like us’, e.g. by conservationists who would like landscape architects to accept the beauty of natural habitats ‘as they are’. As a consequence of this discrepancy in attitudes towards nature, most of the biological information on how green and urban spaces can be made compatible for both human needs and the needs of other species do not penetrate deeply into planning procedures, in particular when they propose to create islands of wilderness in urban areas. Similarly, insightful literature such as ‘gardening for birds’ is mainly read by people who would like their gardens to be similar to nature, rather than to those who would like to design with nature. Thus, in cities, there is a constant battle between ‘design’ and ‘nature’, that is mostly won by design, and rarely by ‘nature’, as landscape architecture is in general more closely aligned with human needs. On the other hand, the preservation of biodiversity and ecosystem services is today one important aim in city planning, and thus requires solutions that go beyond this traditional controversy.

## ANIMAL-AIDED DESIGN AS A METHOD TO MAIN-STREAM BIODIVERSITY INTO URBAN PLANNING PROCEDURES

Overcoming the discrepancy between landscape architecture and conservation can, in our view, only be achieved when both landscape architecture and conservation accept the approach and methods of the other discipline. The rationale underlying AAD is that conservation and landscape architecture can only be successfully combined if biodiversity preservation is integrated into the landscape architectural design process. Thus, AAD aims at good design for humans that benefits animals. We focus on animals, as conservation is mostly about animals; but our approach may also encompass plants, fungi or other groups of organisms. Our method is based on the following premises:

### (I) Target species need to be selected at the beginning of the planning process

Selecting target species before the detailed planning of a building or road construction commences offers the possibility that species requirements are considered in the landscape architectural design. This is in sharp contrast to the current situation, where a completed or advanced design is confronted with the requirement of species that need to be protected. Thus, we suggest to treat the presence of animals as any other requirement or constraint in the design of an open space, such as the layout of a playground for children, an open-air cinema, or the number of benches or parking spaces required. If this is the case, the habitat requirements of species have the same priority in the planning process as do other functions of the open space – not higher, but not lower either. Because the needs of animals thus become an integral part of the planning process, the current time-delay between initial project development and the consideration of biodiversity is avoided (Fig. 2).

**Figure 2:**
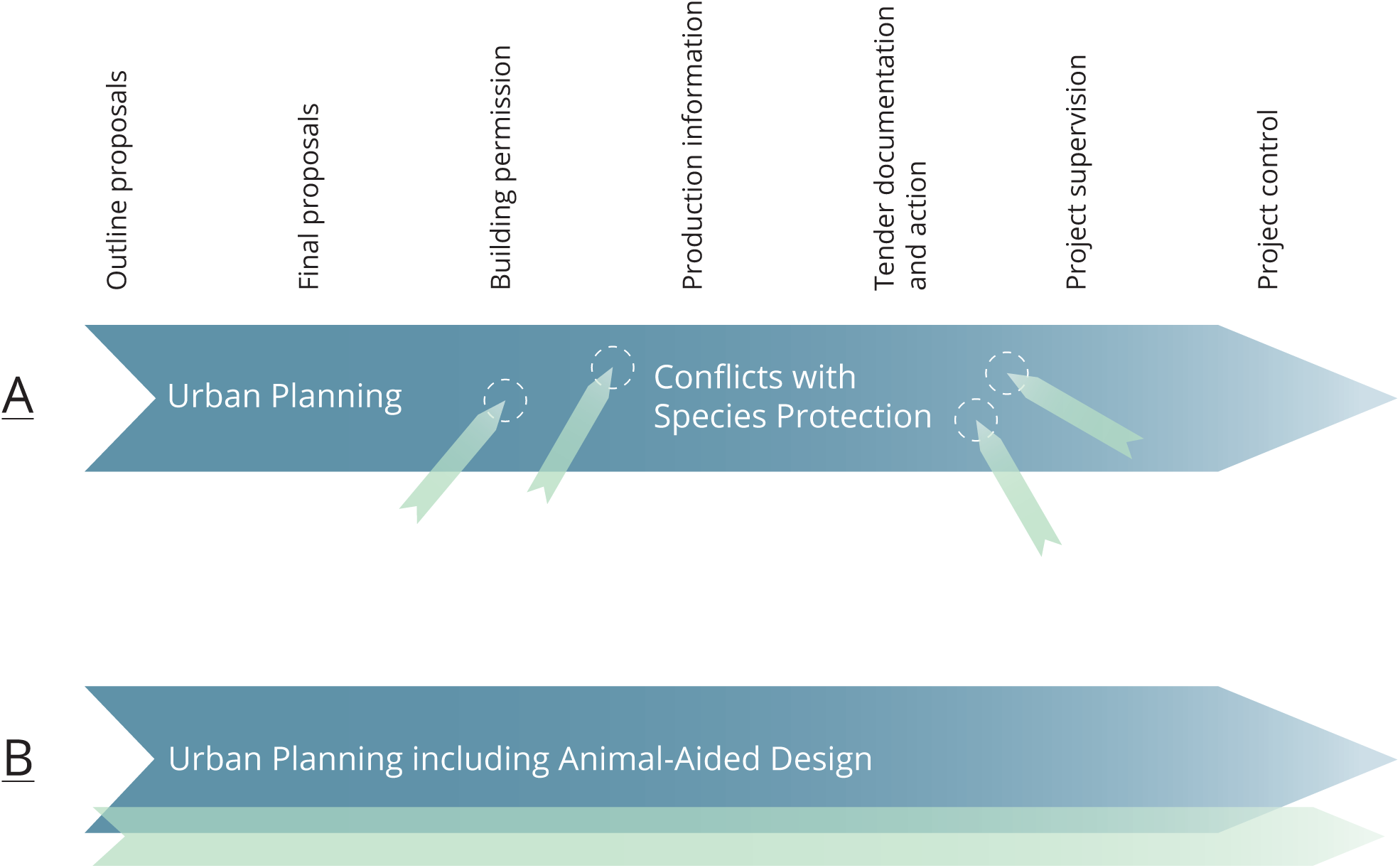
Selecting target species for Animal-Aided Design at the beginning of the planning process avoids the conflict between the design of urban green spaces and conservation (Drawing by T. E. Hauck). A) current situation: a development is planned without reference to biodiversity and the final design is submitted to the authorities for building permission. It is in this advanced stage when it is confronted with the needs of conservation, e.g. through an environmental impact assessment. The presence of a protected species may than require a costly modification of the project (or the removal of the species from the site), a lose-lose situation. B) Animal-Aided Design avoids this conflict by making the habitat requirements of target species to become an integral part of project design, aligning species conservation and the planning process, without delay to the planning process and without surprises.

### (2) Critical needs of the target animals can be identified based on the species life-cycle

The rationale of critical needs is that if the green space provides specific requirements, a viable population of the target species could live in the planning area. AAD thus requires a reductionists approach to the natural history of the species, to identify those elements in the native habitats that are indispensable for the animal. Examples of critical needs are the animal’s food sources, the requirements for nesting sites, or protection from predation, e.g. protection from cats for young birds in the early phase after fledglings have left their nest. The critical needs represent the Hutchinsonian niche of the animal in the open space. In case of a species with large home ranges the designed green space may only make up a small contribution to the habitat where a population lives, but it can become an important part of it.

### (3) The requirements of the animals can inspire the design of the green space

For most critical needs, it does not matter for the animal how they are implemented, in particular how they ‘look’, as long as the solution offered fulfills its functional role. For example, many bird species require a sand bath and a water bath for dusting, drinking and bathing respectively, but there are many examples showing that birds accept a variety of structures acting as sand or water bath (Fig. 3).

**Figure 3:**
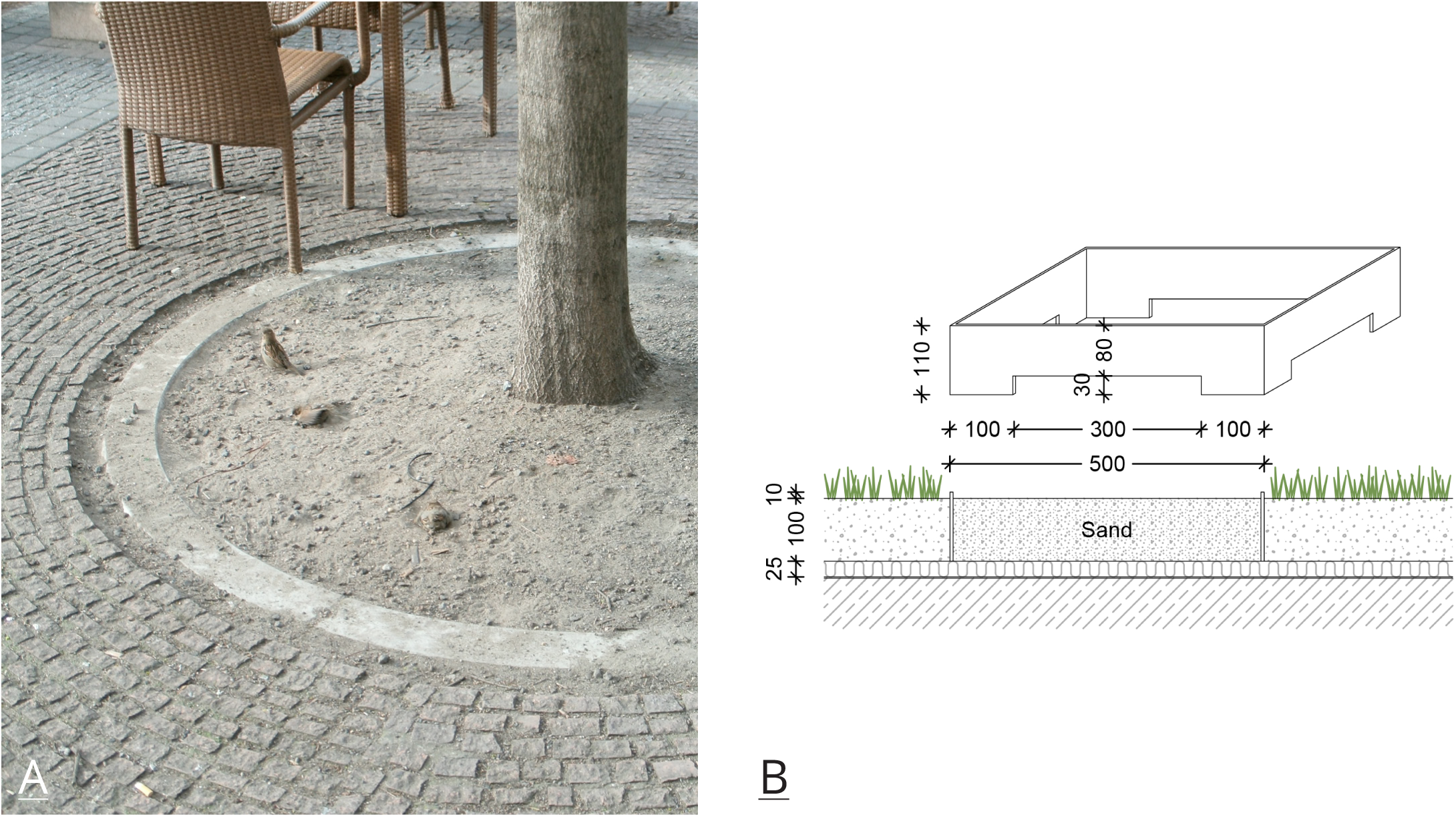
Animals use human-build structures to fulfill their critical needs independent of whether they appear to be ‘natural’ to the human. A) House sparrows *(Passer domesticus)* dusting in one of the few remaining open places with sand underneath trees at Gendarmenmarkt, in the city centre of Berlin, Germany (Photo by W W Weisser). B) Design of artificial sand bath to be placed on a green roof (Drawing and design by Robert Bischer).

In a landscape architectural design of an open space, it is very simple to provide such critical needs. As long as the texture of the sand, or water depth and quality meet the requirements of the animal, it will accept this and the structure will fulfill its function for the animal. Thus, the need to provide a water bath and sand bath is not a constraint for the designer, but may instead inspire the design. This is why we refer to our method as Animal-Aided Design, i.e. design that is enriched, also in its beauty for humans, through including the needs of animals. Importantly, the solutions for the critical needs of animals can and should be multifunctional, i.e. also serving the needs of humans. For example, open water enriches any green space and can be provided by small structures like a small fountain or even an open bowl, thus benefitting both humans and animals such as birds. But this is also true for other needs, a sand bath needed by birds, for example, can be integrated into structures as different as a footpath, a vegetation-free space on a green roof (Fig. 3b) or even a parking space for cars.

## MAKING ANIMAL-AIDED DESIGN WORKABLE FOR THE LANDSCAPE ARCHITECT

Most landscape architects do like animals and welcome it when animals use their open space (as long as they are not undesired species, see below). The biological knowledge required to make an open space suitable for particular species is, however, not easily available for landscape architects. As a consequence, it will not be apparent to a landscape architect, which animals could theoretically be used for AAD at the beginning of the planning process. In addition, even if the landscape architect does choose a target species, e.g. by selecting a species that already occurs in the planning site, it may be difficult to obtain the information on how to adapt the open space to the biology of the species. Consulting a biologist about the habitat requirements of the target species may help, but even a biologist will need to do some research on the habitat requirement of the species, and, for many species, the critical needs will not be absolutely clear, due to lack of knowledge of natural history.

We suggest that to make AAD a workable method, the following is needed:

1) a species portrait listing all critical needs of a particular species (Box 1), and 2) supplementary planning aids that translate the critical needs of species into the design language of landscape architecture.

#### Box 1

Information contained in a species portrait to inform landscape architects and city planners about the biology of the species and its interaction with humans.

- **General characteristics of the species**

- taxonomic affiliation, appearance, geographic range, basic biology
- general habitat characteristics, behavior, natural enemies
- **Human-animal interactions**

- perception of species, e.g. by song/sound
- ecosystem service provisioning
- any interesting behaviour, seasonal and daily times of interaction
- conflicts
- conservation status of species, legal situation
- **Life cycle of the species**

- critical needs of the species ordered by life stage
- planning aids to illustrate how critical needs can be implemented into the design of an urban space (pictograms)
- **More detailed description of life-cycle requirements**

We have developed a format for species portraits that offers both general information on the species and information on the human-animal interface to the landscape architect, i. e. the way in which humans can interact with the animal (Box 1, Fig. 4 and 5). This includes the traits that make a species attractive for humans including the ecosystem services provided by the species. This also includes the aesthetic value of the animal that is shaped by its appearance and behaviours such as singing, mating, nesting or hunting, and, potential conflicts such as how human land use negatively affects the species (e.g. sensitivity to noise), or potential negative effects on humans that need to be avoided (e.g. feces underneath bat entrances in façades). Critical needs of the species are listed for each phase in its life-cycle, from birth to reproduction to death, including courtship, breeding, juvenile and hibernation phases. The planning aids are pictograms graphically illustrating the species requirements (see Figs. 4 and 5 for an example of the common lizard, *Lacerta agilis*).

**Figure 4:**
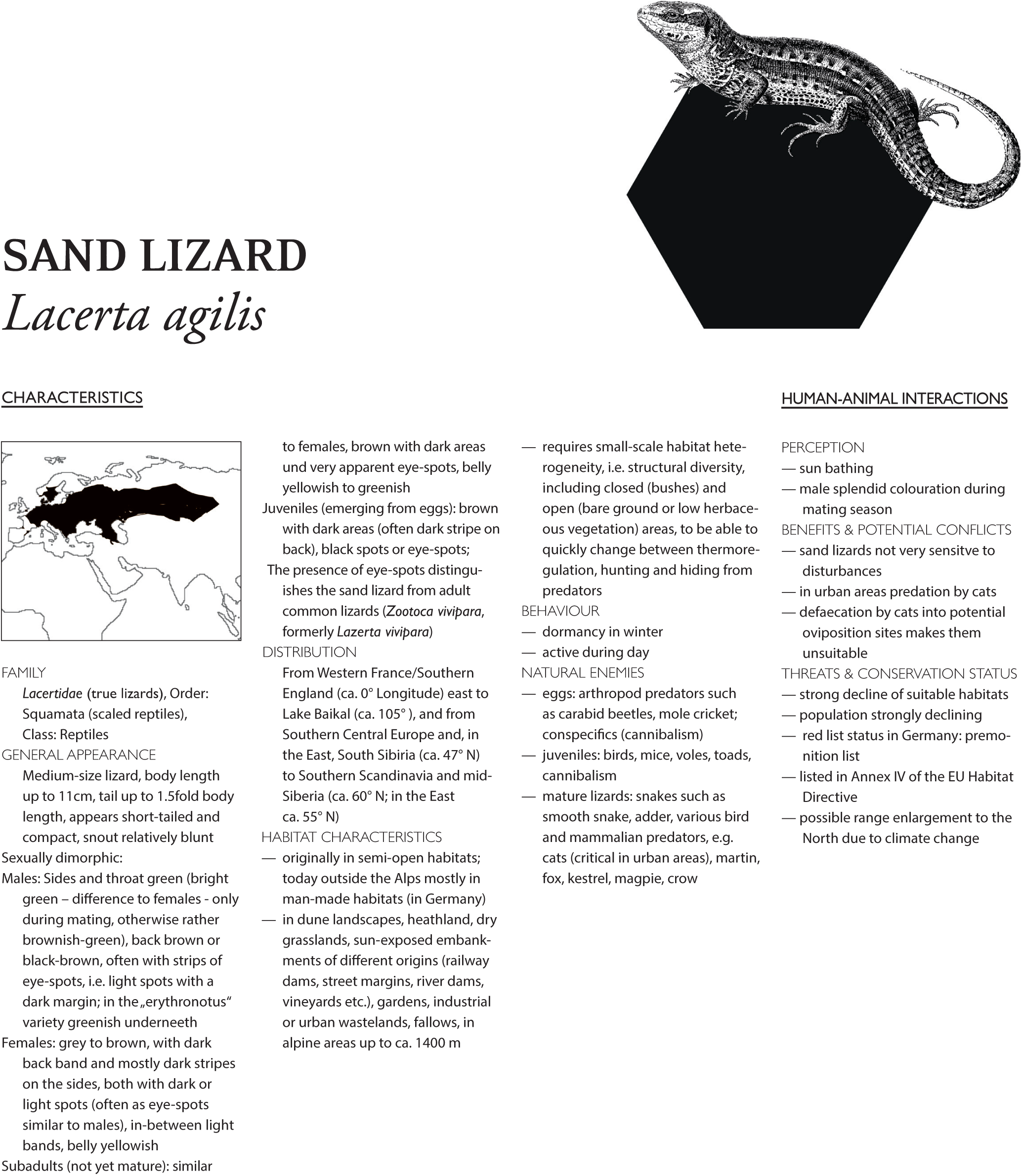
Species portrait of the common lizard *(Lacerta agilis*): General characteristics of the species and human-animal interactions. Not shown is the appendix that lists more detailed information on the biology of the animals relevant for a planner. (Graphic Design by Sophie Jahnke).

**Figure 5:**
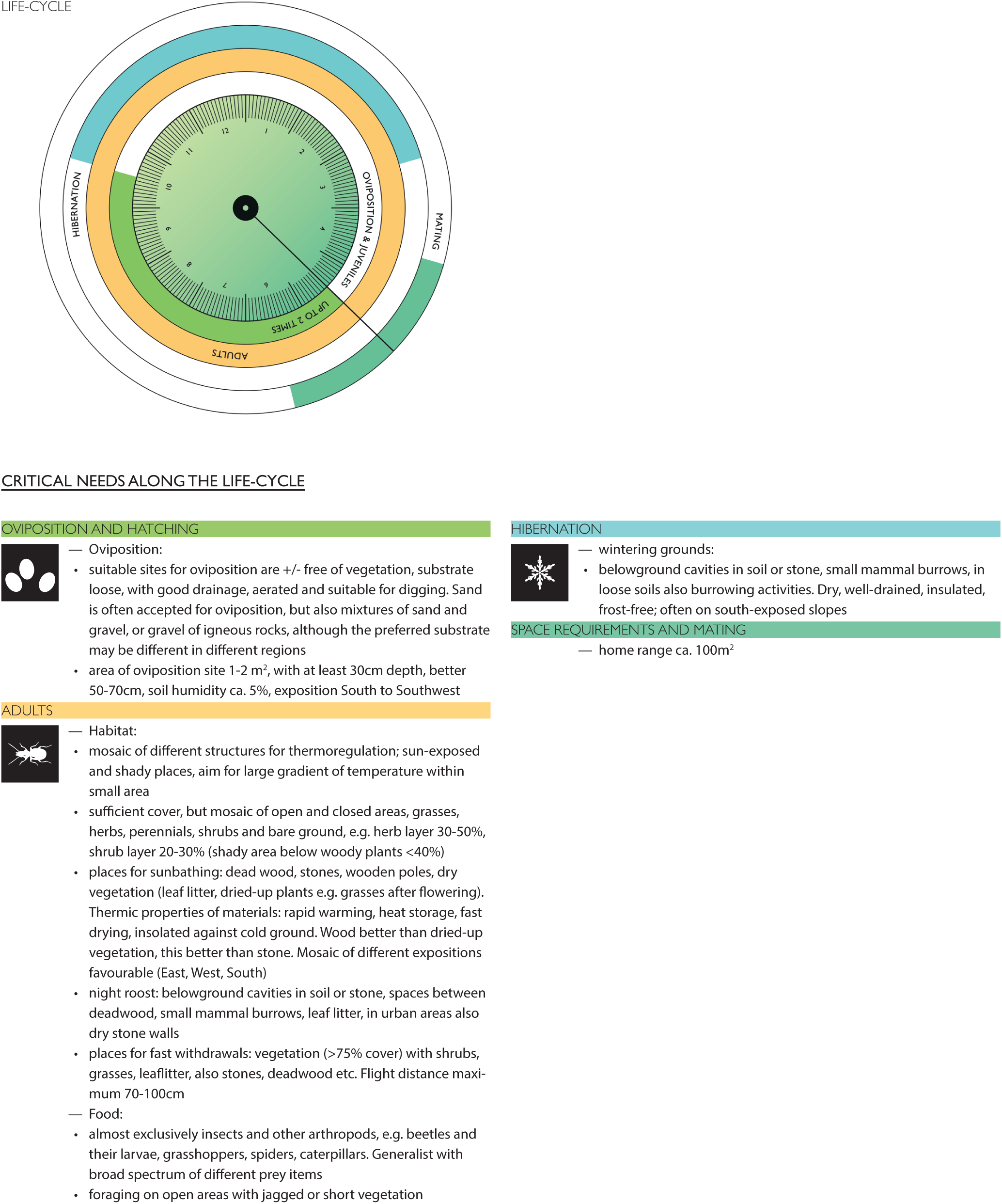
Species portrait of the common lizard *(Lacerta agilis*): Life-cycle of the species and planning aids. (Graphic Design by Sophie Jahnke).

We have chosen the AAD life cycle diagram to be represented by a clock where the hands mark the beginning of the life cycle – birth or eclosion. The different circles in the diagram illustrate the different life phases and the colours illustrate in which months of the year e.g. breeding, takes place. Only if the critical needs of the individuals of a species are fulfilled, the probability of survival and reproduction in the planning area will be high. The landscape architect has to accommodate the needs of the species for every phase of the life cycle into the design. We emphasize this point because this is the weakness of many of the current nature conservation measures such as hanging up nesting boxes or installing bee-hotels. By providing solutions for only a part of the needs – in this case the breeding place, while leaving out other needs –such as food sources – it is left to chance whether the animal will be able to live in the open space. If the design does not cover all critical needs, the plan to establish a species in the project area may fail.

## FROM SPECIES PORTRAITS TO A DESIGN INCLUDING ANIMALS

How should the landscape architect use the species portrait? First, because a species portrait is meant to lists all critical needs of a species, it can be used as a checklist, to test wether a design considers all relevant habitat requirements (Fig. 5).

Second, the planning aids can be used to design both the overall outline of an open space, as well as all necessary details. The pictograms representing the different critical needs can then be implemented into the final design. Importantly, by placing the pictograms into the open space plan, the landscape architect not only demonstrates that a critical need is fulfilled, but also where in the open space it is fulfilled. In the final plan will then be a specific location for each of the critical needs of the animal. This final plan can be used to test not only whether all critical needs have been implemented, but also, whether requirements concerning the relative position of each place are met. For example, during breeding house sparrows will not normally travel further than about 50m, thus all requirements such as the water bath or dusting site must not be further away than 50m from the proposed nesting location (Fig. 6). Similarly, food sources must be in the same circle of 50m.

**Figure 6:**
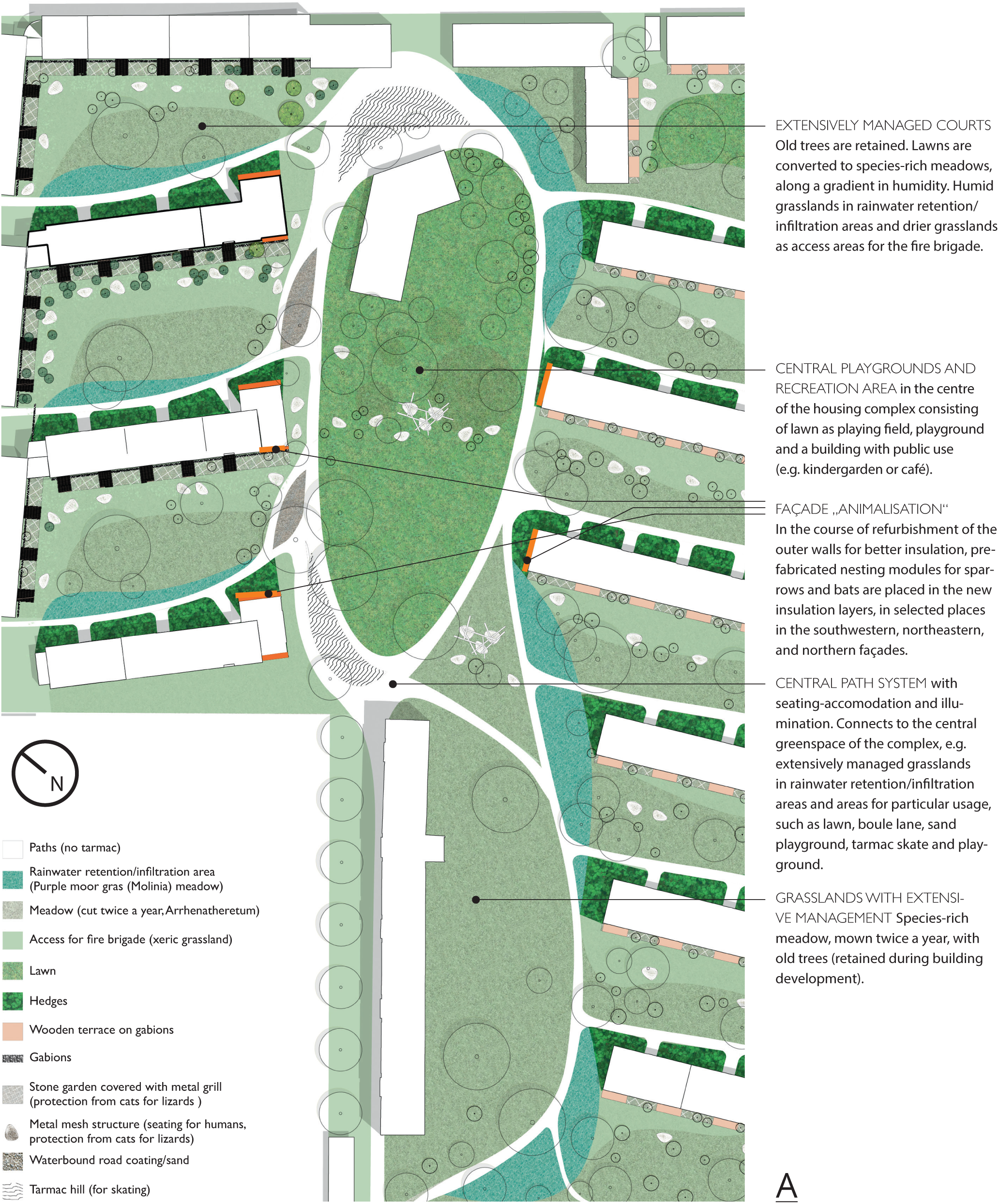
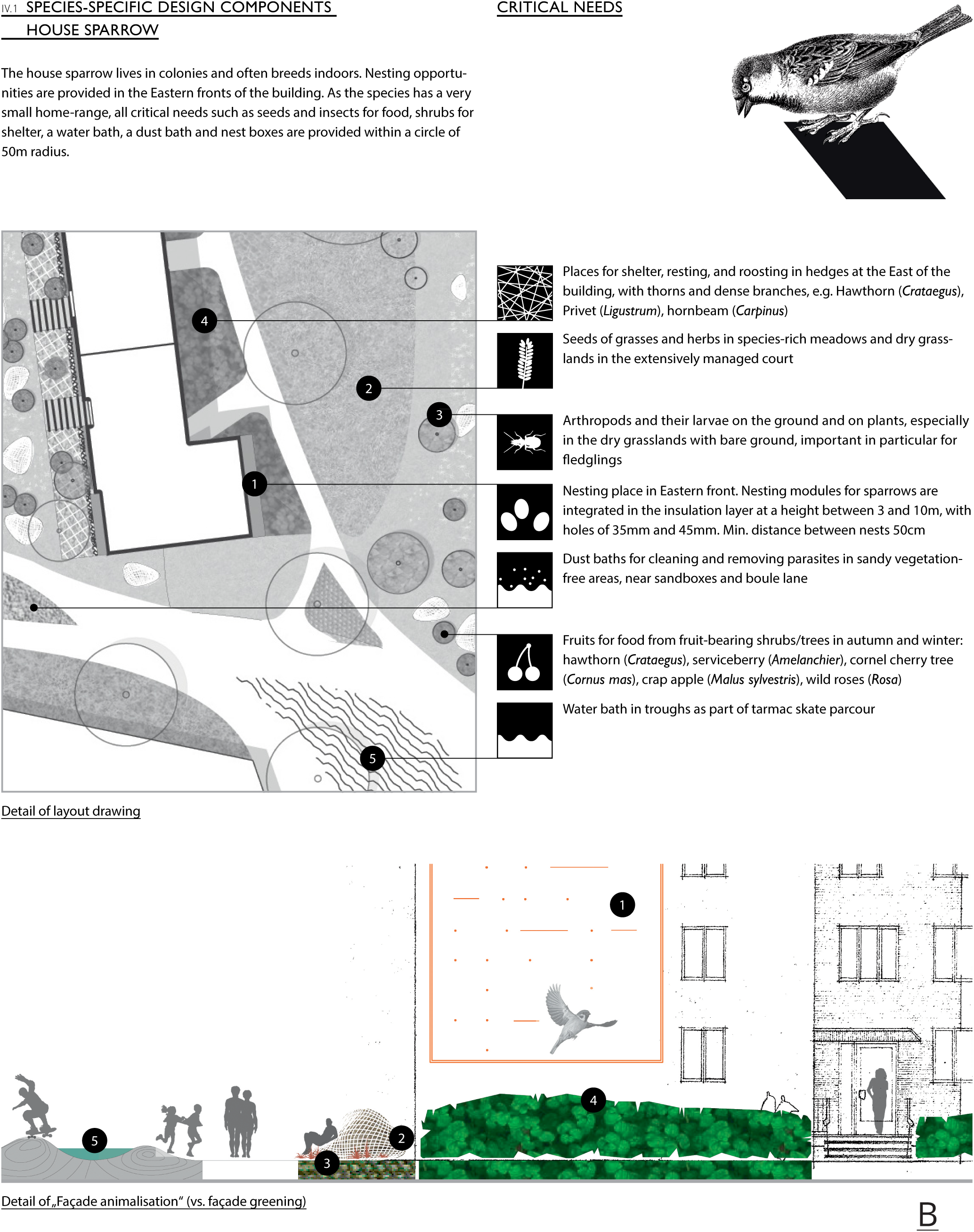
Application of AAD to the refurbishment of an apartment block from the 1960s in Munich, Germany. Such residential areas consisting of several apartment blocks with rental flats were built after the Second World War all over Germany and are now in need of refurbishment including the green space around them. The aim in this theoretical (i.e. not realized) study was to refurbish the building face with increased insulation as an adaptation to climate change, to increase the attractiveness of the greenspace, and to use the public and semi-public space as contribution for the green infrastructure of the city. Target species were the common swift *(Apus apus),* the starling *(Sturnus vulgaris),* the common pipistrelle *(Pippistrellus pippistrellus),* the house sparrow *(Passer domesticus)* and the sand lizard (*Lacerta agilis),* all of which breed in the front (façade) of a building, provided suitable nesting boxes are placed into the outer walls of the building. In the case of the sand lizard a sandy soil suitable for egg-laying can be provided on the ground close to the building. A) Design layout of the central area in the restored residential area. B) Detailed AAD planning for house sparrow where the different pictograms show where in the urban green space particular critical needs of the species are fulfilled. For the house sparrow, all of these needs are fulfilled within 50m of the nest boxes. In the AAD-approach, the planner has to show in the design for the development where a particular species can meet each of the critical needs of the animal. (Design by Rupert Schelle, Georg Hausladen and Sophie Jahnke).

During the design process there will be challenges that need to be overcome. For example, certain structures may potentially be dangerous for humans (e.g. water basins for small children) or the needs of humans and the animal may be incompatible, such as the need of many species for a quiet hibernation place which must not be frequented by humans in winter. It will be the creative part of the planner to find a solution to these challenges, e.g. by spatially separating animal or humans within the development site for some parts of the life-cycle (winter), and joint spaces at other times (summer). Meeting such challenges is, however, nothing special to a landscape architect, as every planning process consists of finding innovative solutions to conflicting demands.

## CHOOSING TARGET SPECIES AND THE VALUE OF AAD FOR CONSERVATION

Two questions generally arise when we present AAD to landscape architects, city planners or ecologists. This first concerns the choice of target species. How to choose a species when planning a building and an open space? We leave the answer to this question deliberately open. This is because, in our view, there may be different reasons why a particular species is chosen for planning a particular open space. An obvious choice are species that are already present at the planning location. These species are likely candidates for AAD when the planners or future inhabitats of a building would like to keep the species and when there is a danger that they will suffer from the development. Thus, pure conservation reasons can be the underlying rationale for choosing a target species. However, we do think that the choice should not be restricted to species that are already there. First, we would like to give the developers the choice to choose species which they consider to be attractive for the design. The song of a blackbird at dusk, the nightly activity of hedgehogs, or the food gathering activities of sparrows during the day, are just a few examples for the many reasons why particular species may be attractive to humans. Second, species absent from the area but of particular conservation concern may also be targets, in order to create new habitats for a species in danger. There may also be additional reasons why a particular target species is chosen.

By allowing target species to be recruited from those not present at the site at the outset and not of (legal) conservation concern, the range of species for which a habitat is provided is vastly extended compared to the current situation, where only species of conservation concern that are already legally protected are subject to conservation measures. Thus, AAD provides the chance to create green infrastructures with animals that would not otherwise be considered. Obviously, there are limits to the choice of species. For example, species that are dangerous to humans, or species that the vast majority humans do not want to have in their vicinity, e.g. the brown rat (*Rattus norvegicus*), will not be suitable target species. Similarly, species whose habitat requirements cannot be met in cities, because they require very large habitats or very specific habitats not present in urban areas, or species that are sensitive to disturbances by humans, are also unsuitable as target species. In the end, the choice of a target species will depend on the outcome of the discussion process among the various stakeholders during project development.

From the conservation point of view, it is an important question whether AAD will only be applied to species such as house sparrows, hedgehogs or robins, e.g. species that are very likable, common in cities and with habitat requirements that are easily met. We think that this must not be the case. In our discussions with planners and developers, interest in a wide range of species has been voiced. Of course it is only when a real building project unfolds that it will become clear what species different stakeholders can agree on, and for what species AAD can be realized, but there is no need for general pessimism. We do think, however, that there is scope for more research on cultural values and conflicts related to wild animals in urban areas, but also on the opportunities and limitations of animal conservation in cities from a biological point of view. There is also need for more research on traits that allow animals to live in cities, as well as more basic research on the natural history of species, to understand their critical habitat requirements.

A second question that we are confronted with is – will it work, i.e. will the species for which the open space is designed, be able to live there?

This is an important question, but one that can only be answered once projects using AAD have been realized. The species portraits that we have already developed have been carefully checked by species experts, but, as with any other conservation measure, the success can only be assessed after implementation. AAD has been applied to a number of test cases, and detailed designs of open spaces have been developed (cf. Fig. 6). Some of these projects are in the process of being realized. Once these and hopefully more projects are completed, it will be possible to judge the contribution of AAD to the creation of green infrastructure.

AAD is a species-centred conservation approach, and its success can be judged using the selected target species. If such species act as umbrella species, many more species in addition to the target species will profit from the design of the open space. Similarly, species that need to be present in an open space to allow for the presence of the target species will also profit, because the design needs to equally include the needs for such species at the site. This could be species needed as food supply, e.g. ants for the green wood-pecker, plants with seeds and insects for the house sparrow or species providing mutualistic relationships, e.g. ants for the target species *Maculinea* butterfly. Thus, AAD may result in providing habitat for a larger range of species, even if the number of target species chosen at the beginning of the project is low. We believe the real contribution of AAD to conservation lies in a) the chance of keeping a species at a site where it would normally be ousted, despite all conservation laws, and b) the chance of creating new habitats where in the current situation this would not be the case.

## CONCLUSIONS

With current rates of urbanization and conservation laws that mostly only succeed to preclude the worst damage to biodiversity in city development, it is necessary to mainstream biodiversity conservation into urban planning strategies. We believe that Animal-Aided Design is a methodology that can help to align the aims of urban planners and conservationists, by making animals an integral part of urban planning strategies. AAD may thus help to overcome the cultural separation into the human world and wildlife that has many repercussions in the way urban development and conservation interact today.

## ACKNOWLEDGEMENTS

We thank the Bavarian State Ministry of the Environment and Consumer Protection for funding (TUF01UF-65043), the GEWOFAG GmbH, Munich, and the Bavarian Society for the Protection of Birds (LBV) for fruitful discussions. We thank Rupert Schelle, Georg Hausladen and Sophie Jahnke for their contribution to the species portrait and the Munich case study presented in this paper; and Anita Schäffer Agnes Wagner, Wolfgang Völkl, Gabriela Jofré and Sylvia Weber for help with collecting the biological information of the sand lizard and the house sparrow, and Christine Jakoby for proofreading.

